# Glycaemia and albumin glycation rates as fitness mediators in the wild: the case of a long-lived bird

**DOI:** 10.1101/2024.07.22.604617

**Authors:** Adrián Moreno-Borrallo, Pierre Bize, Sarahi Jaramillo-Ortiz, Christine Schaeffer, Fabrice Bertile, François Criscuolo

**Affiliations:** University of Strasbourg, CNRS, Institut Pluridisciplinaire Hubert Curien, UMR 7178, 67000 Strasbourg, France; Swiss Ornithological Institute, Sempach, Switzerland; National Proteomics Infrastructure, ProFi, FR2048 Strasbourg, France

**Author notes:** These authors share senior authorship (Fabrice Bertile, François Criscuolo).

**Keywords:** Glucose, albumin glycation, ageing, swifts, glycaemia, fitness

## Abstract

Glucose is a vital metabolic component in the functioning of organisms, but it can also bind to biomolecules through non-enzymatic glycation reactions that result in loss of functions. While the effects of glycation on health have been well demonstrated in biomedical research, little is known about the effects of glycation in wild animals. Here, we studied how plasma glucose levels and albumin glycation rates vary with age and are related to fitness in a relatively hyperglycaemic long-lived bird, the Alpine swift (*Tachymarptis melba*). We measured plasma glucose and albumin glycation levels before and after reproduction in adult females of known age (2-14 years), showing that, while glucose levels increased in parallel with body mass, albumin glycation rates decreased within this period. Albumin glycation, but not glucose, varies with age, peaking at 5 years, consistently with other age-related parameters previously reported in this species. Interestingly, higher plasma glucose levels before reproduction were related to increased fledging success up to a certain threshold. In addition, in terms of dynamics, females gaining more mass lowered more their glycation levels, while those gaining less mass and lowering the more their glycation levels laid more eggs. Finally, higher body mass and plasma glucose levels after reproduction predicted a higher survival probability to the next season, whereas higher albumin glycation predicted lower survival, although in an age-dependent manner. Our study highlights adult plasma glucose and glycated albumin levels as new potential markers of ageing and fitness that should be further explored in this species.

## Introduction

Understanding why some individuals are more performant than others at passing genes to the next generation is one of the main tenets of evolutionary biology. To answer this question, we need not only to measure fitness *via* various indicators of individual survival and/or reproduction, but also to understand the underlying physiological mechanisms regulating organisms’ morphology and performance, and in turn fitness. An interesting physiological mechanism in this respect involves blood glucose and its metabolic by-products resulting from the reaction of sugars with nitrogen compounds such as amino acids like lysine or arginine, namely glycation (Maillard 1912). Although glucose plays an essential role in supplying energy to tissues, its binding to proteins, lipids and nucleic acids through non-enzymatic reactions can also result in a loss of function of the glycated molecules (see e.g. Bakala et al. 2012; Dinda et al. 2015; Suravajjala et al. 2013).

The role of glucose and glycation on the health and performance of organisms is well studied in medical research on laboratory animals and humans. We know that high blood sugar levels are linked to diabetes, a major human disease in developed countries today (Sun et al. 2022). The morbidity related to diabetes is thought to be mediated by several mechanisms, including non-enzymatic glycation (Brownlee 1994). This reaction occurs readily under physiological conditions and can lead to the formation of advanced glycation end-products (AGEs) (Cerami, et al. 1986). AGEs are toxic compounds whose accumulation rate in tissues depends on protein turnover (Verzijl et al. 2000) and metabolic health (Uruska et al. 2019). AGEs accumulate in particular in the case of several pathologies such as cardiovascular or neurodegenerative diseases (see e.g. Poulsen et al. 2013; Chaudhuri et al. 2018; Twarda-clapa et al. 2022; Khalid, et al. 2022). Furthermore, the relationship between glycaemia and fitness is supported by studies in captivity showing that high blood glucose levels and poor glycaemia regulation are linked to mortality rates in zebra finches (Montoya et al. 2018; 2022) and in certain primates, including humans (Palliyaguru et al. 2021), while glucose supplementation can improve survival and immunity in Drosophila (Galenza et al. 2016). In contrast, very little is known about the consequences of glycaemia and glycation in natural populations.

The case of birds is of particular interest given that their glycaemia is the highest within vertebrates, doubling that of mammals (Polakof et al. 2011). However, except from rare cases, birds usually do not show the pathologies (i.e. diabetes) associated with so high glycaemic levels that would be mostly fatal for mammals (partly reviewed in Van de Weyer and Tahas 2024). Protein glycation might therefore constitute a good indicator of health status and a possible fitness predictor. In birds, only three studies have previously investigated the link between haemoglobin glycation and measures of fitness in the wild. They revealed links of haemoglobin glycation with age and survival probability (Récapet et al. 2016) as well as with phenology and fledgling production (Andersson and Gustafsson 1995) in the collared flycatcher (*Ficedula albicolis*) and with chick growth in American kestrels (*Falco sparverius*; Ardia 2006). However, as these studies used mostly non-specific glycation detection methods, more research is needed on the prevalence of glycation in birds and its general relevance as a proxy of fitness.

Unlike previous studies of glycation in wild birds (Miksik and Hodny 1992; Rosa 1993; Andersson and Gustafsson 1995; Beuchat and Chong 1998; Ardia 2006; Récapet et al. 2016; Ling et al. 2020), we assessed here plasma albumin glycation levels instead of glycated haemoglobin levels in erythrocytes. Albumin is the most abundant plasma protein and its glycation levels can be considered as an alternative marker for disease progression monitoring due to its relevance in regulating processes such as blood pressure and oxidative status (Furusyo and Hayashi 2013; Kohzuma et al. 2021). Albumin glycation levels are representative of short-term glycaemia levels (few weeks instead of several months for haemoglobin in humans), so it provides a higher resolution tool for glycaemic regulation (Inaba et al. 2007; Kim and Lee 2012). Importantly, albumin is much more exposed than haemoglobin to blood circulating glucose. Haemoglobin is indeed protected from glucose in erythrocytes given that the transport of glucose inside erythrocytes in birds seems to be virtually non-existent and energy production within erythrocytes depends very little on glycolysis (Johnstone et al. 1998). Accordingly, a previous study shows that, in captive adult zebra finches, plasma albumin glycation levels are rather high whereas no glycation was observed on haemoglobin (Brun et al. 2022).

In this study, we investigated age-related variations in glycaemia and albumin glycation levels in a natural population of a relatively long-lived bird, the Alpine swift (*Tachymarptis melba*; median and maximum lifespan of 7 and 26 years, respectively; Fransson et al. 2023). We also sought to determine if glycaemia and albumin glycation levels are related to fitness, so they can be subjected to natural selection. Finally, we investigated whether glycaemia and albumin glycation could reflect the cost of reproduction, mediating a trade-off between reproductive success and ageing. As this bird species is long-lived, adults are expected, in line with life history theory, to favour maintenance and survival over current reproduction, therefore limiting current effort, as it would be advantageous for maintaining future reproduction prospects (e.g. Sæther et al. 1993). Adult Alpine swifts weigh about 100 g (Dumas et al. 2024) and feed on aerial insects caught exclusively in flight. Given their relatively small size and insect-based diet, their glycaemia is expected to be high, even for birds. Indeed, in birds, blood glucose levels correlate negatively with body mass (Kjeld and Ólafsson 2007; Braun and Sweazea 2008; Tomasek et al. 2019), and animals with high protein intake have higher levels of hepatic gluconeogenesis, thus maintaining sustained high glucose levels relatively independently of their feeding levels (Migliorini et al. 1973; Myers and Klasing 2018).

## Materials and methods

### Species and study colonies

The Alpine swift is a socially monogamous bird that breeds in colonies of a few to hundred pairs in cliffs or in the roof spaces of tall buildings. For this project, data were collected in 2023-2024 in three urban colonies of Alpine swifts located under the roof of clock towers in the Swiss cities of Biel (Stadkirche; about 60 breeding pairs), Solothurn (Bieltor; about 40 breeding pairs) and Luzern (Hofkirche; about 30 breeding pairs). There is an easy access to the nests in these colonies (buildings), which have been monitored for overs 70 years in Solothurn, 40 years in Biel and 10 years in Luzern. Each year, nests are monitored to record the number of eggs laid and hatched as well as the number of chicks fledged (see e.g. Bize et al. 2006; 2008; 2014). Female Alpine swifts produce a single clutch per year of 1 to 4 eggs (modal clutch size is 3). Both parents then share breeding duties, incubating the eggs for 18 days and feeding their offspring until 50 to 70 days after hatching, at which point they fledge. Chicks are ringed at 15 days of age and, given that many individuals are locally recruited (Bize et al. 2017), around 70% of the adult birds in this population have been ringed as chicks and therefore have an exact known age. These long-distant migrants arrive in Switzerland in mid-April, start laying eggs in May and leave in September for their wintering grounds in West and Central Africa (Meier et al. 2020).

### Sample collection and analyses

Alpine swifts were captured under the legal authorisation of the Swiss Federal Agency for Environment, Forests and Landscapes (ringing permit #2235). Blood sampling was performed under a licence of the Veterinary Services of the Cantons Berne, Solothurn and Luzern (National license #34497).

As part of the adult monitoring, each year adults are captured after their arrival from spring migration (late April to early May; i.e. pre-breeding period) and at the end of the reproductive period (August) before they leave for their wintering grounds. All the adults in a given colony are captured on the same day, after dusk, using trap doors that are manually closed after they entered their colony (i.e. building) to roost for the night. After the traps are closed, all the birds are immediately captured by hand and kept in bags until they are measured (for a description of the different measures, see Moullec et al. 2023) and blood samples are taken from some of them. Blood (ca. 150μl) is collected from a toe using heparinized Microvette®(Sarstedt). They are kept on ice before being centrifuged (3500 rpm, 4°C, 10 min), and then the plasma is aliquoted and frozen at -20°C. This processing of blood samples (i.e. centrifugation and -20°C storage) is carried out within 4 hours of collection.

In 2023, the pre- and post-breeding captures of adults took place between May 1 and 4 and between August 9 and 12, respectively, with thus a period of about 100 days separating the two sessions of capture. Due to logistic limitations in the number of samples that could be analysed in the laboratory for glycation levels, we selected samples from 36 females (12 from Biel, 19 from Solothurn, and 5 from Luzern). We restricted our sampling design to one sex to minimize the variance associated with sex, and in turn increase the statistical power associated with analyses on small sample sizes. We chose females because we aimed to study the links with reproduction, and we expected stronger links in females than males (Bize et al 2008). To investigate the links with age, we took care to have a widespread age range (mean ± SE = 6.64 ± 0.57 minimum-maximum = 2 – 14).

Glycation levels were determined using a mass spectrometry-based method, as previously described in Brun et al. 2022. Our measures of albumin glycation rate correspond to the percentage of glycated albumin relative to total plasma albumin. Glucose levels were determined by a Contour Plus (Ascensia® diabetes care) portable glucometer.

We estimated individual annual survival from 2023 to 2024 by looking the presence/absence of the birds in their colony between May and July 2024 following capture sessions at night in spring and during daytime during reproduction. Birds that are not re-seen in 2024 are supposed dead since breeders show no breeding dispersal (Bize et al. 2017), and the probability to recapture a bird that is still alive is virtually 1 in our colonies (Bize et al. 2006).

### Statistics

First, we investigated sources of variation in blood glucose levels, albumin glycation levels and body mass using general linear mixed models where, as fixed explanatory variables, we entered the sampling period (2 factor levels: pre-versus post-breeding), body mass on the night of capture (except for the model on mass itself), and chronological age. When analysing the sources of variation in glucose levels, we also tested for stress effects by including the sampling time in our model. When analysing the sources of variation in albumin glycation levels, we included the measures of glucose levels as explanatory variable to test whether females with higher glycaemia had higher albumin glycation levels.

Besides, we investigated whether pre-breeding glycaemia and glycation levels explained female reproductive performance in the same year by performing a set of models testing the effects of pre-breeding glucose and glycation levels, as well as pre-breeding body mass and chronological age on a series of reproductive traits used as proxies of fitness. In this sense, we analysed, in four different models, effects of the above explanatory variables on: (i) clutch size, (ii) brood size at fledging, (iii) hatching success (proportion of eggs that hatched), (iii) and fledging success (proportion of hatchlings that fledged). This approach allowed testing whether pre-breeding physiological traits may constraint female reproductive performance (see e.g. Metcalfe and Alonso-Álvarez 2010; Stier et al. 2012 for similar discussions with oxidative stress). To test for possible costs of reproduction on the state of physiological traits after breeding (see e.g. Rose and Bradley 1998; Harshman and Zera 2006), we then investigated how post-breeding measures of glycemia and albumin glycation were affected by either clutch size or brood size at fledging (tested in two separated models for each physiological variable, i.e. a total of four models) after controlling for post-breeding mass, chronological age, pre-breeding values of the variable in question (either glucose or glycation) and post-breeding glucose values in the model explaining glycation.

Furthermore, we also investigated how changes in glucose or albumin glycation levels during the breeding seasons (computed as the post-values minus the pre-values) are related to reproductive effort (either clutch size or number of chicks fledged, in two separated models). We also controlled for age and changes in body mass in these models, and for changes in glucose levels in the model testing how changes in glycation levels affected reproduction.

In all the models, we entered the colony identity, as well as female identity (nested within colony) in models with repeated measures from the same females, as random intercept to control for pseudo-replication. To account for possible non-linear effects of age, body mass, glucose levels and sampling time on the response variables, we included both linear and quadratic effects in our starting models using raw polynomials. In models with significant quadratic effects, segmented analyses were performed *a posteriori* to explore the possible existence of a breakpoint (i.e. threshold) separating different linear relationships on either side of this breakpoint. Afterwards, a model including a variable, called “pre_break”, with an assigned value of 0 after the breakpoint and 1 before, and the interaction of such variable with the variable for which the breakpoint was calculated, was fitted. This was repeated swapping the values of the “pre_break” variable to 0 before and 1 after the breakpoint such that both slopes could be estimated. Finally, we performed generalized linear mixed models with a binomial error structure and a logit link function testing the effects of both post-reproductive (August) plasma glucose and albumin glycation, controlling for post-reproductive body mass and age, and the dynamics over the reproductive period (differences between before and after reproduction, i.e. May and August) on survival probability (as the recapture or not of the individual in May 2024). For the models testing the dynamics, body mass differences were included instead of post-reproductive body mass. All the variables included quadratic terms in the initial full models.

In models where glycaemia, glycation or body mass values were entered as covariables, these were centred in order to better interpret their intercepts (Schielzeth 2010). A stepwise backwards procedure eliminating the quadratic components, the hour and/or the mass when these were not significant was used to simplify the models, comparing the AICs and BICs with the *anova* function of R, and selecting the ones with the lowest values. For two individuals sampled in August, we were unable to detect any glycated form of albumin, making them outliers. Therefore, these two measures were excluded from the final models on glycated albumin. We used general linear mixed models with appropriate distribution for our response variables: Gaussian for glucose and glycation levels, Poisson with a logit link for clutch size and brood size at fledging, and binomial for the proportion of eggs that hatched (hatching success) and the proportion of hatched chicks that fledged (fledging success). Models were ran using the *lmer* function of the *lme4* R package, plus *lmerTest* for obtaining p-values (Bates et al. 2015; Kuznetsova et al. 2017). P-values under 0.05 were reported as significant, and between 0.1 and 0.05 as trends.

## Results

### Variation of glycaemia and albumin glycation levels before and after reproduction and with age

Glucose levels were significantly higher after reproduction than before reproduction (mean ± SE glucose levels in mg/dl before versus after reproduction: 328.8 ± 12.3 versus 374.8 ± 7.7; **Table 1**; **Figure 1A**). Glucose levels were not significantly related to body mass or chronological age (**Table 1 A**).

Albumin glycation levels were significantly higher before reproduction when compared to after reproduction (mean ± SE albumin glycation levels before versus after reproduction: 25.39 % ± 1.19 % versus 24.48 % ± 0.38 %; **Table 1B**; **Figure 1B**). Albumin glycation levels linearly decreased with increasing body mass (**Table 1 B**; **Figure 2A**). Albumin glycation levels varied non-linearly with age (**Table 1 B**; **Figure ESM1** and **Figure 2B**). Follow-up analyses using a segmented approach indicated a breakpoint between 5 and 6 years of age, with albumin glycation levels being significantly lower between 2 and 5 years of age than between 6 and 14 (estimate ± SE: 2-5 years 25.22 % ± 1.54 %; 6-14 years: 28.89 % ± 1.14 %; *P* = 0.024). The albumin glycation levels tended to decrease with age after the breakpoint (slope ± SE: -0.156 ± 0.09; *P* = 0.095) and this slope tended to be different to the one from before the breakpoint as showed by the outcome of the interaction term (estimate ± SE: 0.646 ± 0.318; *P* = 0.051), while no significant trend was found for the slope calculated before the break point (slope ± SE: 0.223 ± 0.14; *P* = 0.123). Variation in glycation levels was not explained by variation in plasma glucose levels (**Table 1B**). Finally, body mass increased after reproduction (estimate ± SE: before = 100.6 g ± 2.3; after = 105.41 g ± 0.99; t = 4.87, *P* < 0.001).

**Table 1.**
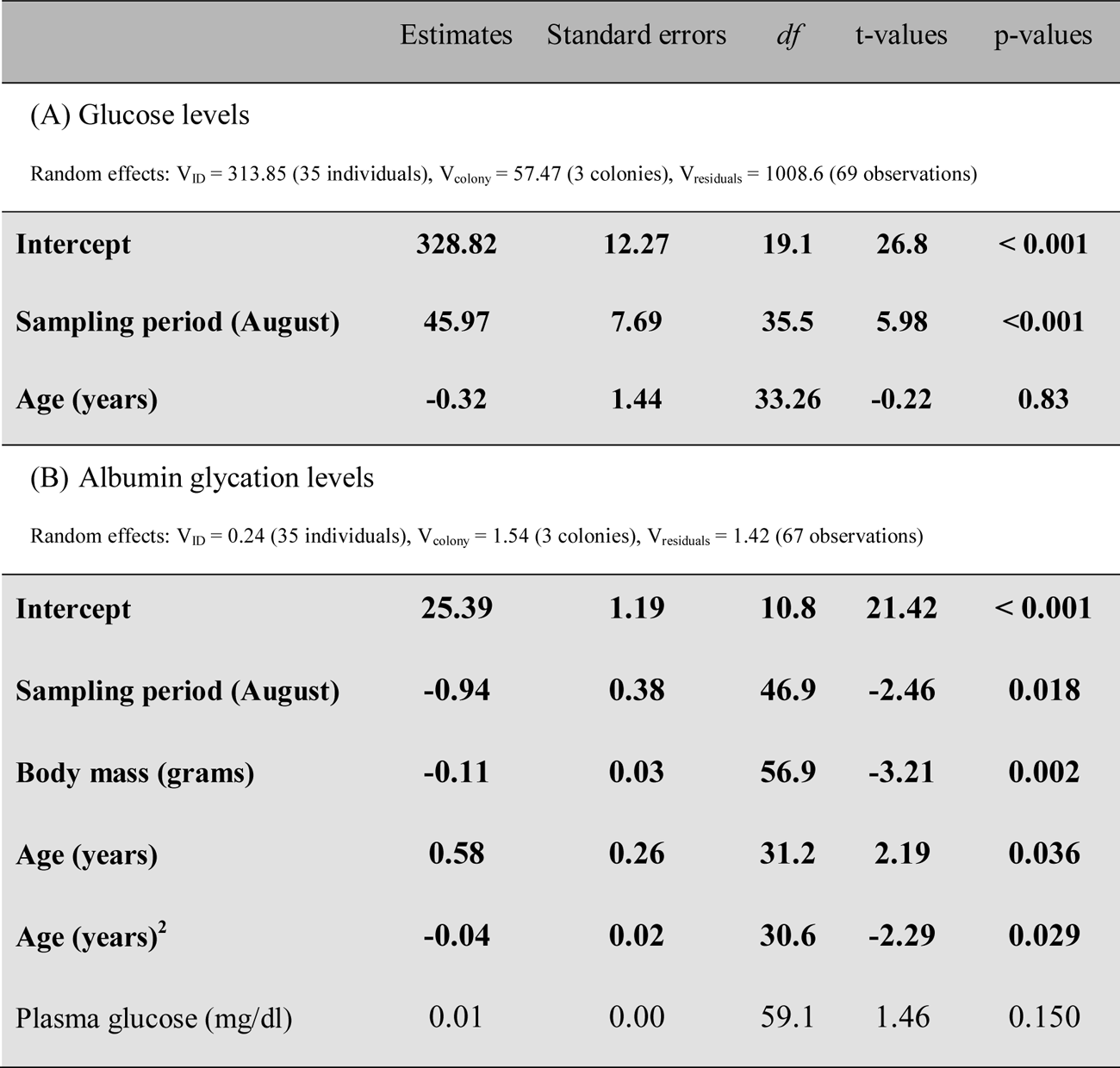
Results of a general linear mixed model showing the effects on glucose. (A) and albumin glycation (as a percentage of total albumin) (B) of the sampling period (before and after reproduction), body mass, age (using a 2-level polynomial approach), and plasma glucose in adult female Alpine swifts. Both glucose and mass are centred to better interpret the intercept. The reference level of the sampling period is before reproduction. Significant predictors are indicated in bold. Bird identity and sampling colony were entered as random effects.

|  | Estimates | Standard errors | df | t-values | p-values |
| --- | --- | --- | --- | --- | --- |
| (A) Glucose levels |  |  |  |  |  |
| Random effects: $V_{ID} = 313.85$ (35 individuals), $V_{colony} = 57.47$ (3 colonies), $V_{residuals} = 1008.6$ (69 observations) | | | | | |
| **Intercept** | **328.82** | **12.27** | **19.1** | **26.8** | **< 0.001** |
| **Sampling period (August)** | **45.97** | **7.69** | **35.5** | **5.98** | **<0.001** |
| **Age (years)** | **-0.32** | **1.44** | **33.26** | **-0.22** | **0.83** |
| (B) Albumin glycation levels |  |  |  |  |  |
| Random effects: $V_{ID} = 0.24$ (35 individuals), $V_{colony} = 1.54$ (3 colonies), $V_{residuals} = 1.42$ (67 observations) | | | | | |
| **Intercept** | **25.39** | **1.19** | **10.8** | **21.42** | **< 0.001** |
| **Sampling period (August)** | **-0.94** | **0.38** | **46.9** | **-2.46** | **0.018** |
| **Body mass (grams)** | **-0.11** | **0.03** | **56.9** | **-3.21** | **0.002** |
| **Age (years)** | **0.58** | **0.26** | **31.2** | **2.19** | **0.036** |
| **Age (years)<sup>2</sup>** | **-0.04** | **0.02** | **30.6** | **-2.29** | **0.029** |
| Plasma glucose (mg/dl) | 0.01 | 0.00 | 59.1 | 1.46 | 0.150 |

**Figure 1.**
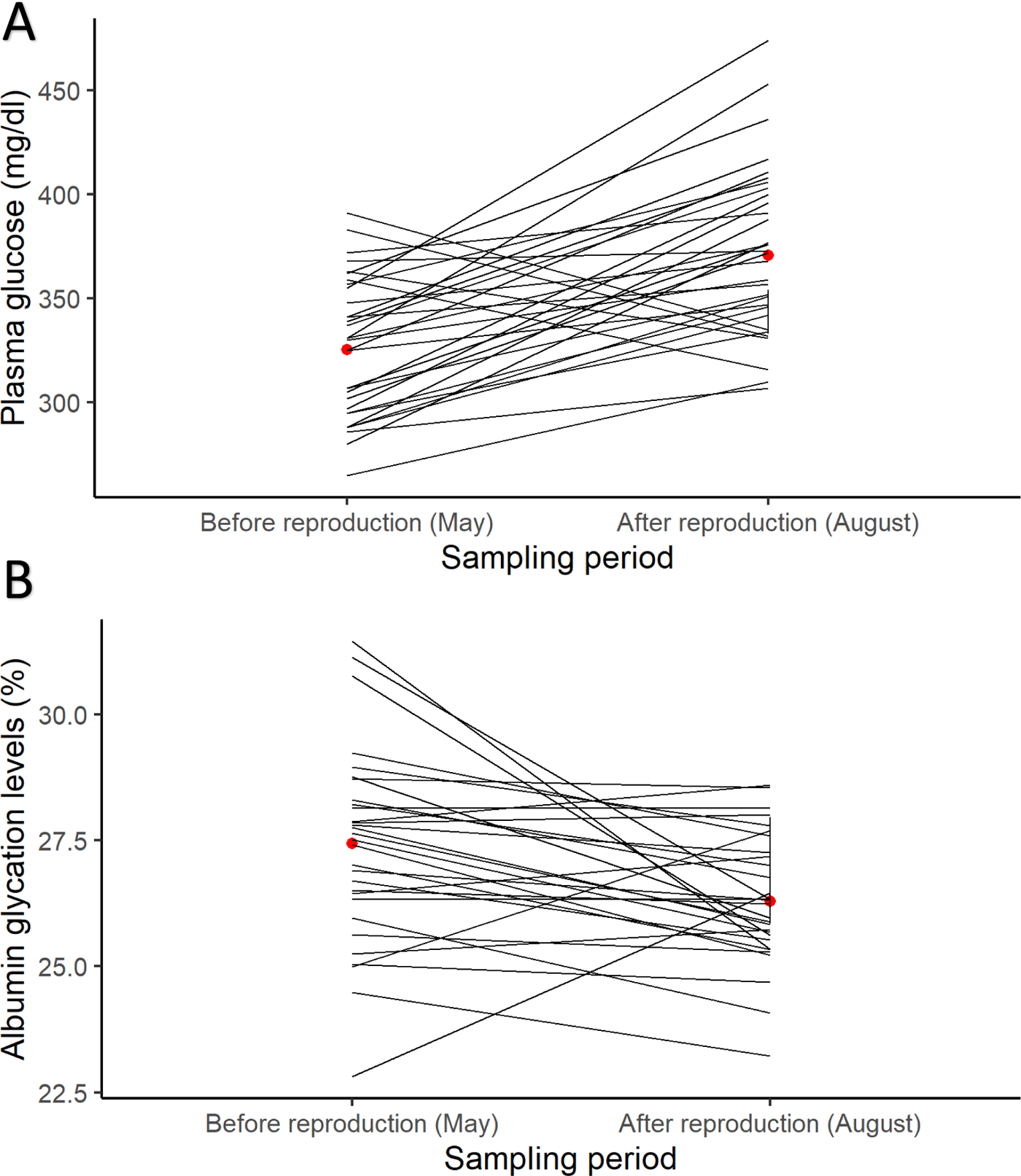
Mean ± SE (A) plasma glucose levels and (B) albumin glycation levels measured before (May) and after (August) reproduction in adult female Alpine swifts (in red). Lines are describing the individual changes between May and August.

**Figure 2.**
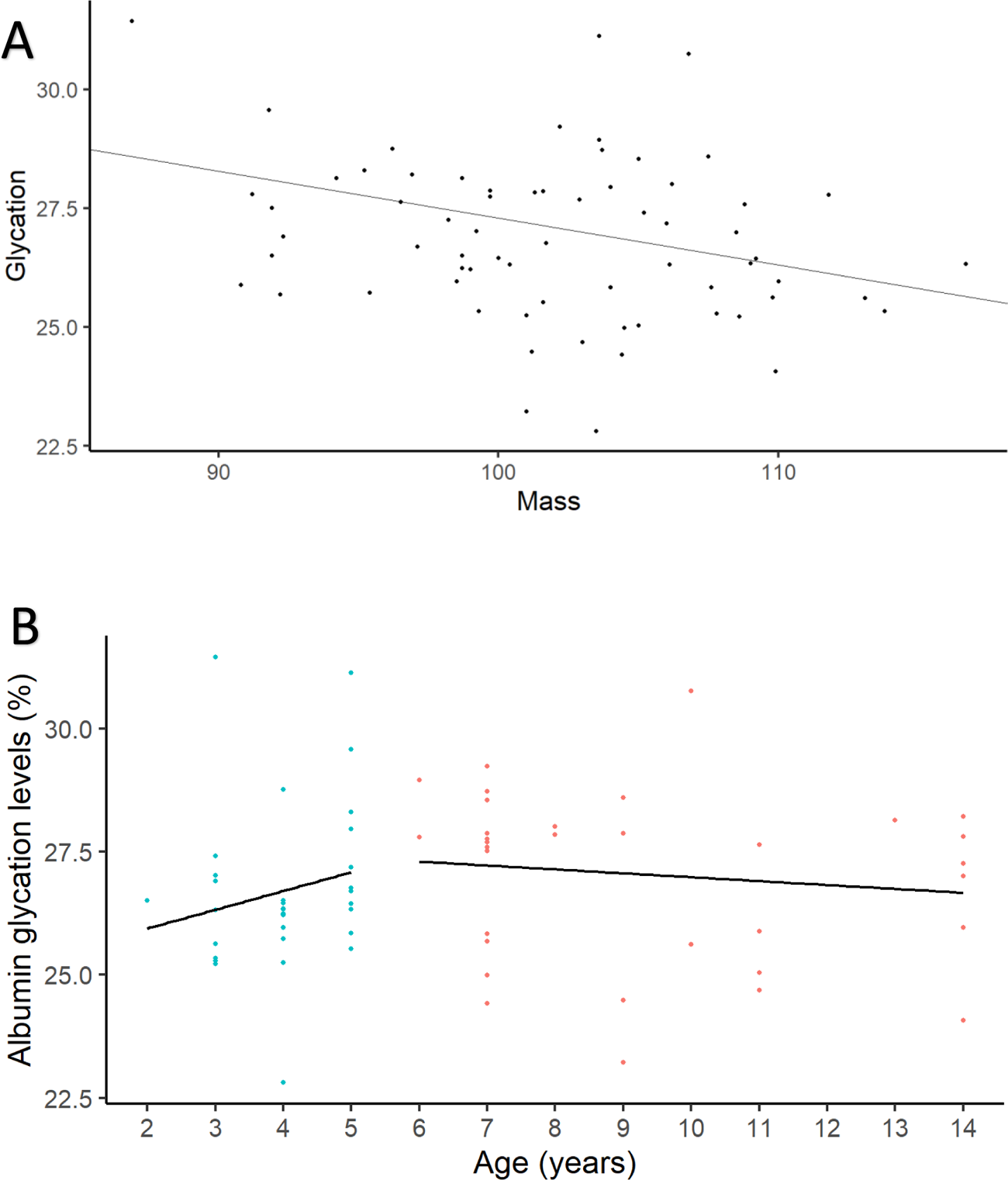
Variation of albumin glycation levels (measured as a percentage of total albumin) in relation to (A) body mass and (B) chronological age in adult female Alpine swifts (following the predictions of a segmented model).

### Glycaemia levels are related to fledging success, but not albumin glycation rates

Plasma glucose and glycation levels measured before reproduction did not significantly explain how many eggs females laid (**Table 2A**) nor how many chicks fledged (**Table 2B**). However, glucose levels, but not albumin glycation, showed a negative trend on hatching success (**Table 2C**), and plasma glucose levels, but not albumin glycation rates, measured before reproduction (May), significantly explained the proportion of chicks fledged from a brood, after controlling for female age, following positive linear and negative quadratic effects (see **Table 2D** and **Figure 3**).

Segmented analyses showed a breakpoint at 307 mg/dl with fledging success showing a trend to be lower below the breakpoint (*P* = 0.095). The fledging success tended to increase with glucose before the breakpoint (estimate ± SE: 0.017 ± 0.009; *P* = 0.074) and this slope tended to be different from that observed after the breakpoint, as shown by the outcome of the interaction term (estimate ± SE: -0.017 ± 0.01; *P* = 0.088). No significant trend was found after the breakpoint (slope ± SE: -0.0008 ± 0.005; *P* = 0.881). Overall, this suggests that there is a limit to plasma glucose levels beyond which fledging success no longer increases as plasma glucose levels rise.

No effect of clutch or brood size at fledging (i.e., reproductive effort) was found on plasma glucose or albumin glycation values measured at the end of the reproduction period (in August), after controlling for glucose/glycation levels measured at the start of the season (in May) (see **Table ESM1.1**).

**Table 2.** Results of a generalized linear mixed model with a logit link function on binomial data of **A** clutch size, **B** brood size at fledging, **C** hatching success, i.e. proportion of eggs that hatched, and **D** fledging success, i.e. proportion of hatchlings that fledged. Age, mass and plasma glucose in May (before reproduction) are included in the model with quadratic effects, and albumin glycation in may only as a linear predictor. Albumin glycation is measured as a proportion of glycated vs total albumin. Significant estimates are indicated in bold.

|  | Estimates | Standard Errors | t-values | p-values |
| --- | --- | --- | --- | --- |
| **(A) Clutch size** |  |  |  |  |
| Random effects: $V_{\text{colony}} = 0.007$ (3 colonies), $V_{\text{residuals}} = 0.201$ (33 observations) | | | | |
| **Intercept** | **0.99** | **0.085** | **11.61** | **<0.001** |
| Glucose before reproduction (mg/dl) | -0.0006 | 0.0009 | -0.655 | 0.512 |
| Albumin glycation before reproduction (%) | -0.001 | 0.02 | -0.064 | 0.949 |
| **(B) Brood size at fledging** |  |  |  |  |
| Random effects: $V_{\text{colony}} = 0$ (3 colonies), $V_{\text{residuals}} = 1.18$ (33 observations) | | | | |
| **Intercept** | **1.424** | **0.189** | **7.53** | **<0.001** |
| Glucose before reproduction (mg/dl) | -0.0007 | 0.006 | -0.126 | 0.9 |
| Albumin glycation before reproduction (%) | 0.074 | 0.103 | 0.716 | 0.479 |
| **(C) Hatching success** |  |  |  |  |
| Random effects: $V_{\text{colony}} = 2.185 \times 10^{-10}$ (3 colonies), $V_{\text{residuals}} = \text{NA}$ (33 observations) | | | | |
| Intercept | 373.55 | 236.87 | 1.58 | 0.115 |
| Glucose before reproduction (mg/dl) | -0.041 | 0.023 | -1.78 | 0.075 |
| Albumin glycation before reproduction (%) | -0.249 | 0.385 | -0.645 | 0.519 |
| Mass (g) | -7.653 | 4.831 | -1.584 | 0.113 |
| Mass<sup>2</sup> (g) | 0.039 | 0.025 | 1.589 | 0.112 |
| Age (years) | 0.587 | 0.333 | 1.762 | 0.078 |
| **(D) Fledging success** |  |  |  |  |
| Random effects: V<sub>colony</sub> = 0.00008 (3 colonies), V<sub>residuals</sub> = NA (28 observations) |  |  |  |  |
| **Intercept** | **-49.90** | **15.42** | **-3.24** | **0.001** |
| **Glucose before reproduction (mg/dl)** | **0.19** | **0.02** | **10.91** | **<0.001** |
| **Glucose<sup>2</sup> before reproduction(mg/dl)** | **-0.00** | **0.00** | **-33.90** | **<0.001** |
| Albumin glycation before reproduction (%) | 0.46 | 0.31 | 1.50 | 0.133 |
| **Age (years)** | **-2.21** | **1.11** | **-2.00** | **0.046** |

**Figure 3.**
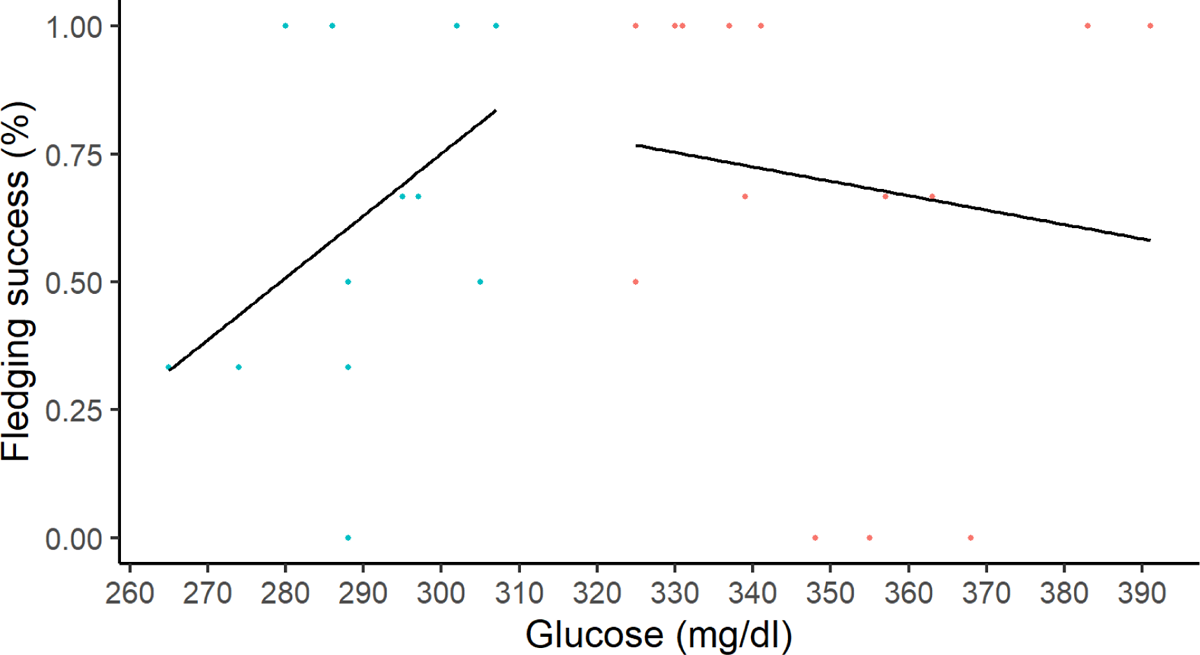
Variation in fledging success measured as the proportion of hatchlings that fledged in function of plasma glucose levels (in mg/dl) measured in May, before reproduction (following the predictions of a segmented model).

### Changes in albumin glycation rates are related to those in body mass and to clutch size

Albumin glycation rate was found to decrease between the start (May) and end (August) of the breeding season, while glucose and body mass increased in parallel over the same period in adult female Alpine swifts (see above). Regarding the dynamics (i.e. differences in the parameters between May and August), we found a significant negative effect of body mass difference on glycation difference (Estimate ± SE =-0.219 ± 0.055; *P* < 0.001, with brood size at fledging as a covariable). In other words, females whose body mass increased the least during the breeding season had a smaller decrease in glycation levels. A similar relationship was obtained when using clutch size as a covariable (estimate ± SE =-0.243 ± 0.061; *P* = 0.0142, see **Table ESM1.2**). When testing the influence of glycation dynamics on clutch size, we found a significant negative effect of body mass and glycation difference on clutch size (**Table 3**). In other words, females with the least reduction in glycation levels laid fewer eggs. Changes in plasma glucose levels were not influenced by any of the breeding output variables (**Table ESM1.1**), nor did they in turn influence them (**Table 3**).

**Table 3.** Results of generalized linear mixed models on (A) clutch size, (B) brood size at fledging, (C) hatching success, i.e. proportion of eggs that hatched, and (D) fledging success, i.e. proportion of hatchlings that fledged. The fixed predictors are the difference between before and after reproduction of plasma glucose in mg/dl, albumin glycation as a percentage of total albumin and mass in grams. The model also includes age in years with a quadratic component. Significant estimates are indicated in bold.

|  | Estimates | Standard Errors | t-values | p-values |
| --- | --- | --- | --- | --- |
| **(A) Clutch size** |  |  |  |  |
| Random effects: $V_{\text{colony}} = 0.013$ (3 colonies), $V_{\text{residuals}} = 0.156$ (31 observations) | | | | |
| **Intercept** | **1.17** | **0.18** | **6.58** | **<0.001** |
| Glucose difference | 0.0005 | 0.0006 | 0.97 | 0.334 |
| **Glycation difference** | **-0.04** | **0.02** | **-2.3** | **0.022** |
| **Mass difference** | **-0.03** | **0.01** | **-3.46** | **0.001** |
| Age (years) | -0.06 | 0.04 | -1.57 | 0.116 |
| Age<sup>2</sup> (years) | 0.004 | 0.002 | 1.83 | 0.067 |
| **(B) Brood size at fledging** |  |  |  |  |
| Random effects: $V_{\text{colony}} = 0.511$ (3 colonies), $V_{\text{residuals}} = 1.04$ (31 observations) | | | | |
| Intercept | 1.66 | 0.51 | 3.25 | 0.065 |
| Glucose difference | -0.001 | 0.004 | -0.23 | 0.816 |
| Glycation difference | -0.191 | 0.314 | -0.44 | 0.542 |
| Mass difference | -0.081 | 0.053 | -1.53 | 0.139 |
| **(C) Hatching success** |  |  |  |  |
| Random effects: $V_{\text{colony}} = 1.652 \times 10^{-10}$ (3 colonies), $V_{\text{residuals}} = \text{NA}$ (31 observations) | | | | |
| Intercept | -0.966 | 1.703 | -0.57 | 0.571 |
| Glucose difference | 0.011 | 0.013 | 0.85 | 0.395 |
| Glycation difference | 0.039 | 0.422 | 0.092 | 0.927 |
| Mass difference | 0.005 | 0.155 | 0.034 | 0.973 |
| Age (years) | 0.447 | 0.317 | 1.41 | 0.158 |
| **(D) Fledging success** |  |  |  |  |
| Random effects: $V_{\text{colony}} = 1.652 \times 10^{-10}$ (3 colonies), $V_{\text{residuals}} = \text{NA}$ (31 observations) | | | | |
| Glucose difference | -0.017 | 0.012 | -1.34 | 0.179 |
| Glycation difference | -0.191 | 0.314 | -0.61 | 0.542 |
| Mass difference | -0.084 | 0.123 | -0.68 | 0.496 |
| Age (years) | -1.725 | 0.929 | -1.86 | 0.063 |
| Age<sup>2</sup> (years) | 0.095 | 0.053 | 1.8 | 0.072 |

The effects on survival probability of plasma glucose, albumin glycation and body mass after reproduction (August), controlling for age, are shown in **Table 4**. While plasma glucose and body mass have linear and quadratic effects on survival (**Figure 4 A, B**) independently of age control, albumin glycation levels only have an effect on survival when age is controlled for (**Figure 4C**).

**Table 4.** Results of generalized linear mixed models on survival to the next season (August 2023 to May 2024) determined by recapture in the colony. Both the complete (A) and the best model by AIC and BIC (B) are shown, to discuss the mediation of age on albumin glycation effects. Significant estimates are indicated in bold.

|  | Estimate | Standard Error | z value | P-value |
| --- | --- | --- | --- | --- |
| *(A) Complete model* |  |  |  |  |
| Random effects: Vcolony = 1.336*10<sup>-17</sup> (3 colonies), Vresiduals = NA (34 observations) |  |  |  |  |
| Intercept | 0.19 | 0.199 | 0.953 | 0.34063 |
| **Glucose after (mg/dl)** | **0.046** | **0.011** | **4.36** | **<0.001** |
| **Glucose<sup>2</sup> after (mg/dl)** | **-0.00005** | **0.000005** | **-9.177** | **<0.001** |
| **Albumin glycation (%)** | **-3.43** | **1.325** | **-2.592** | **0.00955** |
| **Albumin glycation<sup>2</sup> (%)** | **0.059** | **0.025** | **2.394** | **0.017** |
| **Mass (g)** | **0.313** | **0.073** | **4.316** | **<0.001** |
| **Mass<sup>2</sup> (g)** | **-0.001** | **0.0001** | **-8.587** | **<0.001** |
| Age (years) | -0.276 | 0.614 | -0.45 | 0.653 |
| Age<sup>2</sup> (years) | 0.016 | 0.036 | 0.451 | 0.652 |
| *(A) Best model* |  |  |  |  |
| Random effects: Vcolony = 0 (3 colonies), Vresiduals = NA (34 observations) |  |  |  |  |
| Intercept | -12.57 | 19.91 | -0.631 | 0.528 |
| **Glucose after (mg/dl)** | **0.041** | **0.0102** | **4.03** | **<0.001** |
| **Glucose<sup>2</sup> after (mg/dl)** | **-0.00004** | **0.000005** | **-8.15** | **<0.001** |
| Albumin glycation (%) | -1.54 | 1.325 | -1.16 | 0.244 |
| Albumin glycation<sup>2</sup> (%) | 0.022 | 0.025 | 0.912 | 0.362 |
| **Mass (g)** | **0.455** | **0.071** | **6.37** | **<0.001** |
| **Mass<sup>2</sup> (g)** | **-0.002** | **0.0001** | **-14.3** | **<0.001** |

**Figure 4.**
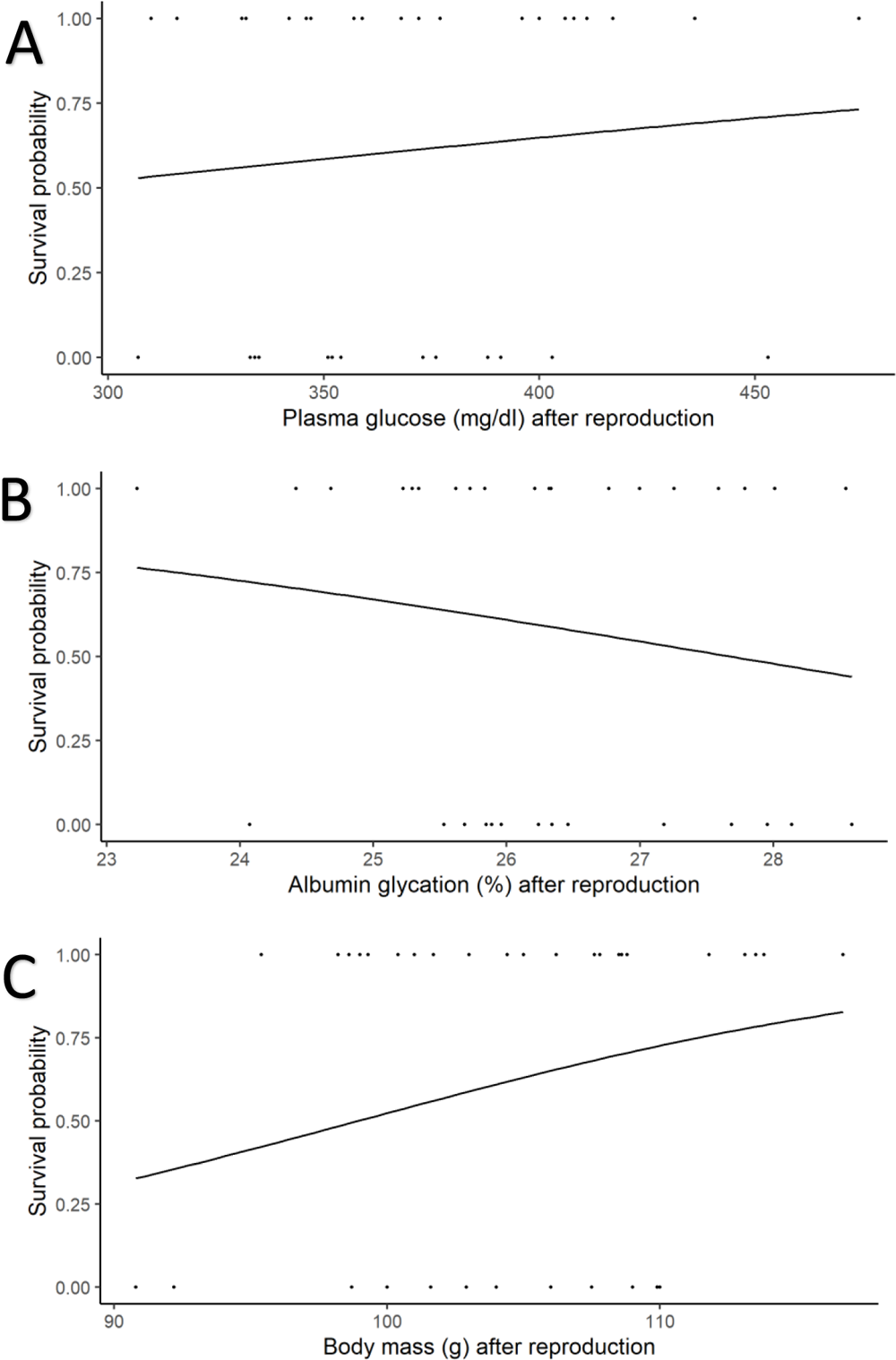
Probability of survival from August 2023 to May 2024 depending on post-reproduction (August 2023) values of (A) plasma glucose levels, (B) albumin glycation rate and (C) body mass.

There was no significant effect of changes in plasma glucose levels, albumin glycation and body mass between pre- and post-breeding stage on the probability of survival (see **Table ESM1.3**).

## Discussion

### Body mass, glycaemia and albumin glycation: related signs of body condition?

Female Alpine swifts had lower levels of plasma glucose and higher levels of glycated albumin before reproduction (May) than after reproduction (August). Alpine swifts are trans-Saharan migratory birds that arrive in Switzerland in April, reproduce between May and August, and then leave in September to return to their wintering grounds in Western Africa (Meier et al. 2020). Hence, blood parameters measured in May could partially mirror a cost of migration, although measures before the migratory event should be performed to discriminate such an effect. Blood glucose levels have been found to positively correlate with residual body mass (adjusted for structural size) in American house finches, *Haemorhous mexicanus*, (Mcgraw et al. 2020), breeding pale-bellied Tyrant-Manakins, *Neopelma pallescens* (Azeredo et al. 2016) and with body mass in garden warblers (*Sylvia borin*) (Jenni-Eiermann and Jenni 1994) and barn swallows (Bendová 2020, but see Lill (2011) and Remage-Healey and Romero (2000)) for contrasted results). Although plasma glucose and body mass were not correlated in our study, they both increased over the course of the reproductive season. This is in contrast with the decrease in mass along the season found by Dumas et al. (2024) in the same Alpine swift population; however, that study focused on body mass measurements during the breeding season, excluding the pre- and post-breeding measurements reported in the present analysis. An increase in feeding frequency of the chicks from the pre-laying to the fledging phase could explain the higher glucose levels (Jackson et al. 2023). However, it remains unclear if the observed increase in body mass across the breeding season depends more on muscle size (proteins), as it would be expected when activity levels are increased (Marsh 1984 and discussion in Jackson et al. 2023), or on fat deposition, potentially derived from sugars. Measures of changes in body composition during breeding, apart from merely body mass, would also be fruitful for understanding how dynamics of fuel usage and sparing affect the value of plasma metabolites such as glucose.

The concomitant increase in body mass and glucose and reduction in glycation between May and August may indicate that glycated albumin plasma levels may be a proxy of individual quality (see Wilson and Nussey 2010) in breeding female swifts. This new individual quality marker would therefore add to the various blood metabolites already described to date as markers of quality in birds (Jenni-eiermann and Jenni 1994; Minias and Kaczmarek 2013; Azeredo et al. 2016; Jackson et al. 2023). Individuals of higher quality may therefore be better protected against glycation, either by avoiding it or by being more efficient in quickly clearing glycated proteins from the body. Therefore, restoring an adult physiological status promoting survival would also limit the deleterious effects of glycation over time, and could therefore promote longevity. Interestingly, albumin glycation and, more generally, fructosamine levels (a marker of general plasma protein glycation) are negatively related to BMI (Body Mass Index) in humans (Selvin et al. 2018). A similar result was found here in swifts, i.e. a negative relationship between albumin glycation levels and body mass. Beyond a simple covariation of body mass and glycation, how body mass may relate to glycation levels is not yet well-known, but it is hypothesized to be related with an increase in protein turnover associated to the proinflammatory state induced by an augmented adiposity (see e.g. Chagnac et al. 2003; Koga et al. 2007).

To our knowledge, few studies described glycation status in wild breeding birds. The only publication in this regard found no significant variation in the levels of circulating glucose and fructosamine after breeding in common eiders (*Somateria mollissima*), despite a decrease in body mass (Ma et al. 2020). This discrepancy may have several explanations. Common eiders are capital breeders and they rely mostly on internal lipid stores while fasting during breeding (Parker and Holm 1990). The relationship between plasma glucose, glycation levels and reproduction may therefore not be as straightforward as in income breeders like swifts.

With regard to body mass change in the incubating sex during breeding, we can observe clear differences between bird species, depending mainly on their reproductive strategy (Moreno 1989), with capital breeders losing weight during incubation while income breeders maintain or even increase their body mass, and undergo the main weight loss shortly after hatching. Cox and Cresswell (2014) found that species whose body mass increases the most during breeding also have higher survival rates. They proposed that the lower food predictability under less pronounced seasonality triggers greater adult investment in survival than in reproduction during the breeding season. Swifts forage exclusively on flying insects, whose availability depends on weather conditions (Grüebler et al. 2008), with rainy days severely limiting adult food intake and affecting their body mass (Dumas et al. 2024). Swifts are relatively long-lived birds, and should therefore invest more in their survival (maintaining body mass and body condition) during the breeding season than in current reproduction (see e.g. Charlesworth 1980; Saether 1988), although it would also depend on certain variables, including their pre-breeding condition, which can modulate their resolution of such trade-off (Erikstad et al. 1998). Indeed, Martins and Wright (1993) showed that common swifts (*Apus apus*) rapidly restore their body mass at the end of the nestling phase, when the chicks’ feeding requirements are reduced, thus limiting the cost of reproduction. In our study, we found that body mass increased from laying initiation to fledging stages, which supports the idea of prioritizing investment in adult self-maintenance for breeding individuals.

### Age-related variation of glycated albumin

Albumin glycation in female swifts increases in early life up to the age of 5, before decreasing slightly with age. This typical bell-shape pattern (see e.g. Forslund and Pärt 1995; Saraux and Chiaradia 2022) of albumin glycation levels in relation to age is similar to what has been described before in this species for morphometric and breeding traits (Moullec et al. 2023), as well as for physiological traits such as females’ erythrocyte membrane resistance to oxidative stress (Bize et al. 2008). Moreover, albumin glycation, together with other glycation markers, has been reported to increase with age in humans, although linearly instead of following a bell-shaped pattern (Selvin et al. 2018). Glucose levels, however, did not vary with age, in contrast to what is known from biomedical research in humans and other model species, where we can see for example an increase with age in primates and a bell-shaped pattern in laboratory mice (Palliyaguru et al. 2021). Our data shows also a higher survival of individuals with higher glucose levels, which together with an increased fledging success, suggests our population of swifts is being selected for higher glucose levels, being potentially quite resistant to its pervasive effects. Still, this could also generate an antagonistic pleiotropy affecting ageing (Williams 1957), given the effects of glycation on survival (see below).

Our results on age-related variation of glycated albumin are derived from a cross-sectional study where individuals of different ages are sampled once rather than from a longitudinal study where the same individuals are sampled multiple times at various ages. Findings from cross-sectional studies are therefore incorporating both effects taking place at the individual level (improvement/maturation early in life and senescence late in life) and demographic effects such as the selective appearance and disappearance of phenotypes in the population (Forslund and Pärt 1995; van de Pol and Verhulst 2006). Hence, the increase in glycation levels before the age of 5 could be explained either by an increase in glycation rates at individual level with age or by the late recruitment into the breeding population of individuals with higher glycation rates (i.e. selective appearance: a later age at reproduction in females with a higher glycation level early in life). Female Alpine swifts usually start breeding for the first time between 2 and 5 years of age (Tettamanti et al. 2012). Similarly, a decrease in older age may occur either at an individual level or as a result of the selective disappearance from the population of individuals with higher glycation rates. An intra-individual decrease in glycation levels with age towards the end of life would indicate ‘improved health’, which contrasts sharply with the results of biomedical research in humans, which show an intra-individual increase in glycation levels with age, indicating ‘senescence’ (Selvin et al. 2018). There is considerable support for the hypothesis of selective disappearance in other species. Indeed, glycation of albumin and haemoglobin has been linked to mortality in humans (Wu et al. 2021; Rooney et al. 2022), whereas adult collared flycatchers with higher haemoglobin glycation levels (measured using a human blood kit) were more likely to disappear from the wild (Récapet et al. 2016). Furthermore, our data showed interesting effects of albumin glycation levels on survival to the next season, which seems to indicate that individuals with lower glycation levels have a higher probability of survival, but only when accounting for age, even when age itself does not predict survival. This can be so because the effects of age on mortality rates (potentially senescence) are mediated by factors such as glycation rates or body mass, for which our data seem to show selection in favour of higher body masses. This is in partial disagreement with what Dumas et al. 2024 found, i.e. that such an effect only occurs for non-breeders, whereas we report it for breeders. Nevertheless, they showed this effect for both sexes together, whereas we do it for females alone. This may be relevant as Dumas et al. (2024) also showed different kinds of selection pressures (although through fledging success, not through survival as in our case) between sexes (i.e. stabilizing selection for females and disruptive selection for males) for breeding body mass (i.e. measured in June).

As a conclusion, to finally determine whether glucose and glycated albumin can be reliably used as markers of senescence in swifts, and potentially in other birds, longer longitudinal studies are still needed to determine if the effects are sustained across different years, potentially with contrasting environmental conditions modulating the selective pressures.

### Pre-reproduction plasma glucose levels and albumin glycation dynamics affect female reproductive performance

We found that plasma glucose levels before the start of reproduction were the only factor that significantly predicted fledging success. Hence, body condition of the parents (here evaluated as body mass) did not positively modulate fledging success in swifts, contrary to what has been highlighted before in this species (Dumas et al. 2024) and in other birds (with body mass residuals on size, e.g. Chastel et al. 1995; Moe et al. 2002). This discrepancy in our results may be due to our small sample size, which makes it difficult to establish significant relationships if the effect is subtle. The relationship between pre-reproduction glucose levels and fledging success was quadratic, and a further segmented analysis suggested that starting breeding with low glycaemia is not optimal for reproduction, and a glucose level over a point (estimated to be 307 mg/dl in our model) does not improve reproductive success anymore. Low glycaemia may indicate poor female condition, with deleterious consequences such as impaired adult foraging performance and seldom chick provisioning, two important determinants of fledging success (e.g. Jenni-Eiermann and Jenni 1994, Saraux and Chiaradia 2022). Although this is not an obvious effect according to our data, which seem to indicate a plateau for fledging success after a threshold in plasma glucose (i.e. 307 mg/dl), high glycaemia could also impair the ability of females to rear chicks successfully. This could involve the glycation process for example, although our data did not suggest that albumin glycation levels directly influence reproduction. In contrast, Borger (2024) found that glycated haemoglobin in female zebra finches was positively related with clutch size and offspring production, but as they only measured it after reproduction, costs cannot be separated from constraints. It may thus be concluded that their results indicate that higher egg production and perhaps clutch care induce higher levels of protein glycation, costs we did not find. However, direct comparison between our results and those of Borger (2024) is difficult, as discrepancies may be explained by physiological differences between species studied (Bize, et al. 2014), the context (captive versus wild individuals), the nature of the targeted glycated proteins, and even the analytical method used (non-specific kits versus accurate mass-spectrometry). A more detailed study of glycaemia dynamics during the successive phases of the breeding season (mating, laying, incubation, brooding, fledging) in our species, as previously conducted in others (Remage-Healey and Romero 2000, Gayathri, et al. 2004, Azeredo, et al. 2016, Bendová 2020), would help to better understand how fledging success is affected by circulating glucose levels in the parents.

Interestingly, the decrease in glycated albumin levels during the breeding season was positively related to clutch size in our study, without being reflected in fledging success, suggesting explanations not related to breeding performance but rather to more basic metabolic processes. For instance, our results can reflect a general increase in anabolism during the reproductive period. Reduced glycation levels may be attributed to faster protein (albumin) turnover rates, thereby reducing damage levels (Wada, et al. 2016). This is consistent with the negative relationship between variation in body mass and variation in albumin glycation levels that we recorded during the course of the breeding season: the greater the increase in body mass after reproduction, the greater the reduction in glycation levels. The negative relationship between the number of eggs produced and the increase in body mass probably reflects a trade-off between somatic and reproductive investments (see Williams 2005 for a review of the costs associated with egg laying), while the positive relationship between glycation reduction rate and clutch size might indicate a closer relationship between albumin turnover and egg production. In fact, albumin production should be increased during egg-laying due to its transfer to the eggs (Patterson et al. 1962).

## Conclusions and perspectives

We show here that glucose and glycated albumin levels increase and decrease respectively during reproduction in Alpine swifts, which is paralleled by an increase in body mass. We did not find any costs related to reproductive effort, evaluated using clutch or brood size. Our results on survival show that glycation levels may represent a more important constraint for other phases of the annual cycle, such as the long-distance migration swifts perform. Future studies should explore if changes in glycation levels such as those detected (around 1%) are paralleled by a significant change in other parameters better known to reflect changes in health or fitness. This would help to better understand the underlying mechanisms mediating glycation and fitness. In addition, we established that albumin glycation levels vary with age in a similar way as other age-related parameters in this species, with a peak at 5.5 years, suggesting that glycation may be part of the physiological mechanisms underlying the senescence process. Nevertheless, this is a cross-sectional study, so longitudinal data would be needed to determine if this variation is really linked to ageing per se and not to other demographical phenomena like selective appearance or disappearance, as our results on survival suggest. Finally, glucose levels before reproduction showed a positive effect on fledging success up to a certain threshold, after which a plateau is reached. A negative trend on hatching success was also found, while glycated albumin levels are negatively linked to body mass, and its dynamics to changes in body mass and clutch size, suggesting that they could influence female reproductive fitness. Although the exact mechanisms underlying our results remain unclear, we hypothesize that albumin glycation dynamics may be mainly influenced by protein turnover rate and may thus be representing the general rate of anabolism, and that glucose is likely to be an indicator of parents’ nutritional state, acting positively on chicks’ survival to fledging. To prove these hypotheses, more precise measures of energy metabolites and their dynamics on both parents and offspring, together with body condition and body composition changes during rearing should be determined, to see which nutrients are the most important to determine breeding success and how the costs and benefits of differential allocation between parents and chick survival are shaped.

## Supporting information

ESM1

## Acknowledgements

This research was performed under the funding of an ANR (AGEs – ANR21-CE02-0009). We thank the numerous researchers and fieldworkers who helped at collecting the long-term data and samples used in this publication.

## Authors contributions

FC and FB conceived the original collaboration, set up the project, contributed to the development of the idea and participated in the discussion of the results. AMB contributed to the development of the idea, participated in the field work and sample collection with PB, and made the statistical analyses. CS and SJO performed the glycation measurements by mass spectrometry. PB provided data from the long-term monitoring of the animals, contributed to the sample collection and participated in the discussion of the results. AMB wrote the original draft, later edited by FC, FB and PB. All the authors approved the final draft.

## Statements and Declarations

Authors declare having no competing interests affecting the content of this publication. Data will be made publicly available on Figshare after manuscript acceptance.

