## Supplementary material for "Glycaemia and albumin glycation rates as fitness mediators in the wild: the case of a long-lived bird": ESM1

* These authors share senior authorship.

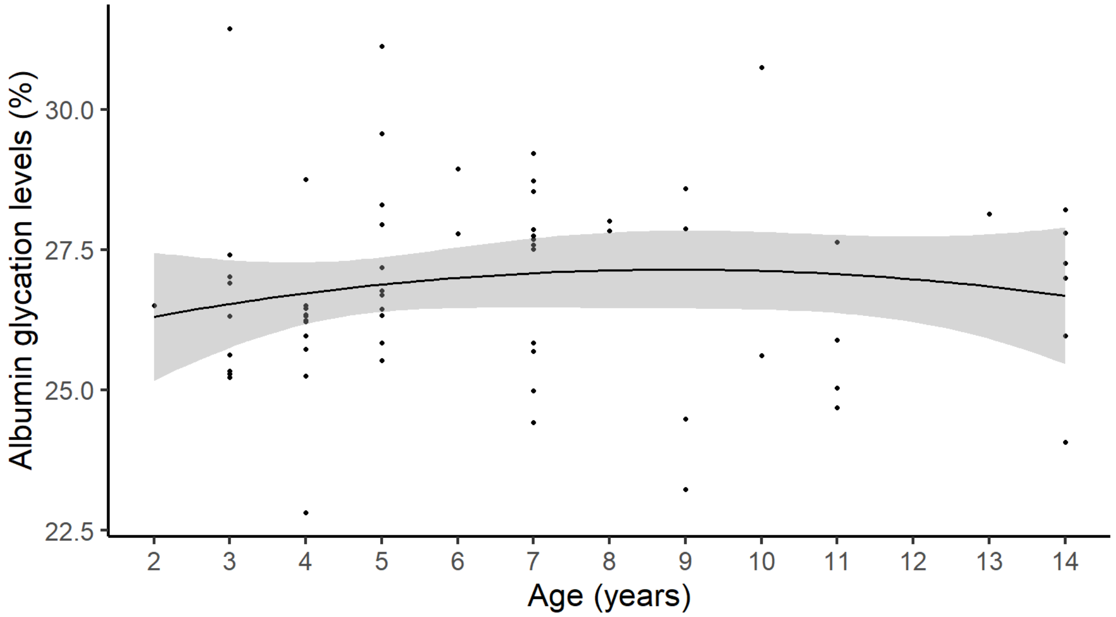


**Figure ESM1.** Variation of albumin glycation levels (measured as a percentage of total albumin) in relation to chronological age in adult female Alpine swifts.

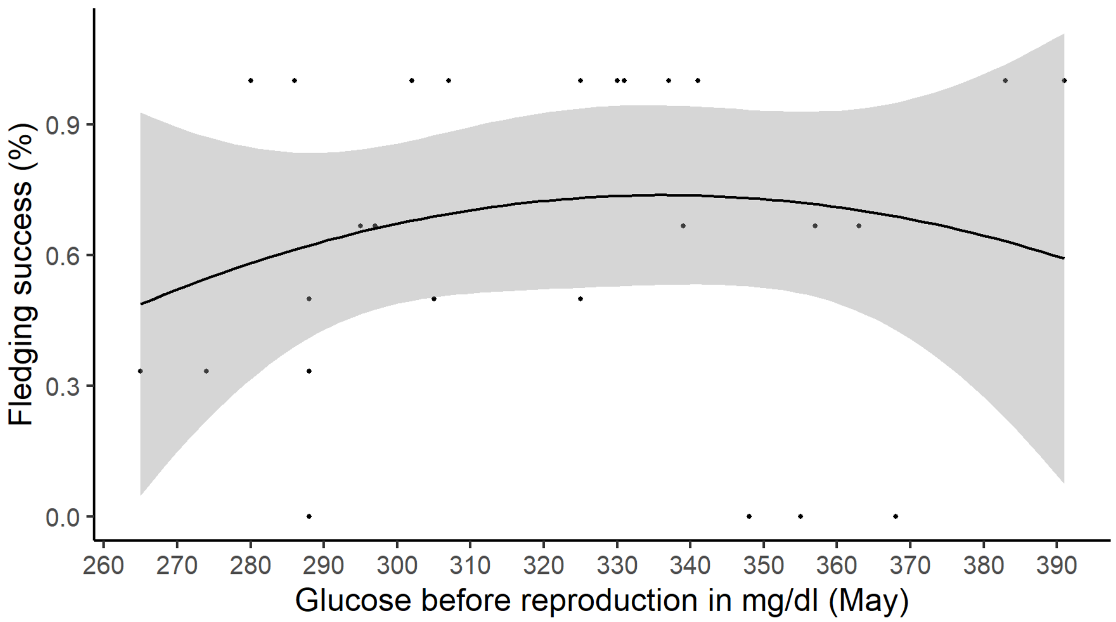


**Figure ESM2.** Variation of fledging success (measured as a percentage of chicks hatched that fledged) in relation to plasma glucose values before reproduction in adult female Alpine swifts.

**Table ESM1.1.** Results of general linear mixed models on the effects of reproduction investment on plasma glucose levels (in mg/dl) difference between before and after reproduction. Two different estimates of reproductive investment (A) clutch size, and (B) brood size at fledging, were considered in different models. The difference in mass in grams is included as a fixed predictor.

| | Estimates | Standard Errors | df | t-values | p-values |
|---|---|---|---|---|---|
| (A) *Clutch size* | | | | | |
| Intercept | 40.83 | 49.45 | 7.26 | 0.826 | 0.435 |
| Clutch size | -1.42 | 17.01 | 11.89 | -0.084 | 0.935 |
| Mass difference | 1.53 | 1.65 | 7.18 | 0.923 | 0.386 |
| (B) *Brood size at fledging* | | | | | |
| Intercept | 40.34 | 18.45 | 3.94 | 2.187 | 0.095 |
| Brood size at fledging | -1.96 | 7.95 | 27.72 | -0.246 | 0.807 |
| Mass difference | 1.45 | 1.65 | 5.96 | 0.879 | 0.414 |

**Table ESM1.2.** Results of general linear mixed models on the effects of reproduction investment on albumin glycation levels (in percentage of total albumin) difference between before and after reproduction. Two different estimates of reproductive investment (A) clutch size, and (B) brood size at fledging, were considered in different models. The difference in mass in grams and the difference in glucose in mg/dl are included as fixed predictors.

| | Estimates | Standard Errors | df | t-values | p-values |
|---|---|---|---|---|---|
| (A) *Clutch size* | | | | | |
| Intercept | 1.5 | 1.79 | 6.14 | 0.837 | 0.434 |
| Clutch size | -0.72 | 0.606 | 10.53 | -1.19 | 0.261 |
| **Mass difference** | **-0.243** | **0.061** | **4.31** | **-3.97** | **0.014** |
| Glucose difference | 0.008 | 0.006 | 26.97 | 1.3 | 0.204 |
| (B) *Brood size at fledging* | | | | | |
| Intercept | -0.5 | 0.63 | 27 | -0.797 | 0.432 |
| Brood size at fledging | -0.028 | 0.28 | 27 | -0.099 | 0.922 |
| **Mass difference** | **-0.219** | **0.052** | **27** | **-4.3** | **0.0002** |
| Glucose difference | 0.009 | 0.006 | 27 | 1.4 | 0.174 |

**Table ESM1.3.** Results of general linear mixed model on the effects of difference between before and after reproduction in plasma glucose (in mg/dl) and albumin glycation levels (in percentage of total albumin) on survival to the next season (August 2023 to May 2024). The model shown is the best after model step back-wise selection process with AIC and BIC is performed.

| | Estimates | Standard Error | z value | p-values |
|---|---|---|---|---|
| Random effects: Vcolony = 0 (3 colonies), Vresiduals = NA (31 observations) | | | | |
| Intercept | -0.099 | 0.597 | -0.166 | 0.868 |
| Glucose difference | 0.001 | 0.008 | 0.119 | 0.905 |
| Glycation difference | -0.368 | 0.232 | -1.585 | 0.113 |
